# Disinfection of SARS-CoV-2 contaminated surfaces of personal items with UVC-LED disinfection boxes

**DOI:** 10.1101/2021.03.03.433725

**Authors:** M. Bormann, M. Alt, L. Schipper, L. van de Sand, M. Otte, T. L. Meister, U. Dittmer, O. Witzke, E. Steinmann, A. Krawczyk

## Abstract

The severe acute respiratory syndrome coronavirus 2 (SARS-CoV-2) is transmitted from person-to-person by close contact, small aerosol respiratory droplets and potentially via contact with contaminated surfaces. Here, we investigated the effectiveness of commercial UVC-LED disinfection boxes in inactivating SARS-CoV-2 contaminated surfaces of personal items. We contaminated glass, metal and plastic samples representing the surfaces of personal items such as smartphones, coins or credit cards with SARS-CoV-2 formulated in an organic matrix mimicking human respiratory secretions. For disinfection, the samples were placed at different distances from UVC emitting LEDs inside commercial UVC-LED disinfection boxes and irradiated for different time periods (up to 10 minutes). High viral loads of SARS-CoV-2 were effectively inactivated on all surfaces after 3 minutes of irradiation. Even 10 seconds of UVC-exposure strongly reduced viral loads. Thus, UVC-LED boxes proved to be an effective method for disinfecting SARS-CoV-2 contaminated surfaces that are typically found on personal items.

## Main Manuscript

### Background

Since the end of 2019, a novel coronavirus called the severe acute respiratory syndrome coronavirus 2 (SARS-CoV-2) has been spreading worldwide, thereby causing a major public health issue (*1*). Coronavirus disease 2019 (COVID-19) is caused by SARS-CoV-2 and characterized by symptoms ranging from mild respiratory illness to severe life-threatening pneumonia and acute respiratory distress syndrome (ARDS) (*2*). Reducing the transmission of the virus by suitable preventive measures is highly important to control the pandemic. SARS-CoV-2 is transmitted by direct contact with infected individuals, by virus-containing aerosols and potentially via virus-contaminated surfaces (*3*). Recent studies have shown that SARS-CoV-2 can persist on smooth surfaces such as glass, metal and plastic for up to seven days at room temperature, remaining a potential risk of infection (*4*).

UV-irradiation is an environmentally friendly method to disinfect surfaces from bacteria, fungi and viruses such as SARS-CoV-2 (*5, 6*). Under laboratory conditions, high viral loads of SARS-CoV-2 in cell culture medium could be completely inactivated by UVC-irradiation after 9 minutes of irradiation with a UVC dose of 1048 mJ/cm^2^ (*7*). For private use, commercial UVC-LED boxes are available for the disinfection of personal items such as smartphones, keys, coins or credit cards. However, the performance of such devices on the inactivation of SARS-CoV-2 on surfaces has not yet been investigated. Therefore, in the present study we investigated the ability of two UVC-LED boxes to disinfect surfaces typically found on personal items such as glass, metal and plastics from contamination with high viral loads of SARS-CoV-2.

## Methods

### Cells and Viruses

Vero E6 cells (American Type Culture Collection, ATCC, CRL-1586, Rockville, MD) were cultured in DMEM supplemented with 10% (v/v) fetal calf serum, Penicillin (100 IU/mL) and Streptomycin (100 µg/mL). SARS-CoV-2 was isolated from a nasopharyngeal swab of a patient hospitalized due to COVID-19 at the Department of Infectious Diseases of the University Hospital Essen in April 2020 (*7*). Viral titers were determined by endpoint dilution according to Spearman and Kärber (*8*) and calculated as TCID_50_ (tissue culture infectious dose, 50%).

### Measurement of the emitted light intensity

Two UVC-LED boxes were investigated for their capability to inactivate SARS-CoV-2 (UVC-LED box 1, Horcol; UVC-LED box 2, expondo GmbH, Berlin, Germany). A radiometrically calibrated spectrometer (STS-UV-L-50-400-SMA, Ocean Optics B.V., Ostfildern, Germany) with a sensitivity range between 190 and 650 nm (1.5 nm resolution) equipped with a CC-3-UV-S corrector (Ocean Optics B.V., Ostfildern, Germany) was used to determine the light intensity emitted by the LEDs inside the UVC-LED boxes. The emitted light intensity was determined between a wavelength of 250 and 280 nm at specific distances from the light source (LED) of the UVC-LED boxes. The corrector was placed at the same locations as the virus carriers (UVC-LED box 1: 1 cm and 5 cm horizontal distance; UVC-LED box 2: 1 cm vertical distance). For UVC-LED box 2, emitted light could only be measured at a distance of 1 cm. A measurement at 4 cm distance would have required drilling a hole into the bottom of the box, which was not possible without damaging the box. The data were recorded using OceanView 2.0 Software and visualized using GraphPad Prism 9 (GraphPad Software, San Diego, CA).

### UVC-LED decontamination of SARS-CoV-2 contaminated surfaces

SARS-CoV-2 working stocks (5 × 10^6^ TCID_50_/mL for UVC-LED box 1; 2 × 10^6^ TCID_50_/mL for UVC-LED box 2) were diluted in a defined organic matrix mimicking respiratory secretions (9). In brief, 900 µL of the respective virus stock was added to 100 µL organic matrix consisting of 2.5 mg/mL mucin type I-S, 7.8 mg/mL BSA Fraction V and 11 mg/mL yeast extract (all Sigma-Aldrich, Darmstadt, Germany). Before inoculating the respective virus suspension to glass, metal and plastic carriers, the carriers were sterilized with a UV-lamp emitting 1940 µW/cm^2^ UVC at 254 nm (UV-4 S/L, Herolab, Wiesloch, Germany) (*7*). After sterilizing, 50 µL of the respective virus suspension were placed on the center of the carriers and allowed to dry for 1 h at room temperature. The carriers were positioned at different distances from the UVC-LEDs inside the UVC-LED boxes (UVC-LED box 1: 1 cm and 5 cm horizontal distance; UVC-LED box 2: 1 cm and 4 cm vertical distance). The carriers were irradiated for specific durations (0 s, 10 s, 30 s, 1 min, 3 min and 10 min). Subsequently, the infectious virus was recovered by vortexing the carriers placed in plastic containers (SARSTEDT, Nümbrecht, Germany) filled with 2 mL DMEM for 1 minute. As control, the virus was recovered 10 minutes after drying without irradiation. The experiments were conducted in triplicates and the viral loads were determined by endpoint dilution according to Spearman and Kärber. The means, standard deviations of the viral titers and 90% effective concentration (EC_90_) values were calculated with GraphPad Prism 9 (GraphPad Software, San Diego, CA). The statistical significances were determined with the t-test. Comparisons were considered significant at *P < 0.05; **P < 0.01; and ***P < 0.001.

### Results

SARS-CoV-2 can potentially be transmitted via virus-contaminated surfaces of personal items such as smartphones, keys, coins or credit cards that have been contaminated with the virus. We investigated the performance of two commercially available UVC-LED boxes for virus inactivation on surfaces typically found on personal belongings such as glass, metal and plastic. We used two different UVC-LED boxes for sterilizing, one with lateral UVC-LEDs (Fig. 1A) and one with UVC-LEDs incorporated in the lid of the box (Fig. 1B). Additionally, a mirror was installed to the bottom of the chamber. For UVC-LED box 1, the emitted light intensity was determined with 245 µW/cm^2^ at horizontal distance of 1 cm and 65 µW/cm^2^ in the center of the box at horizontal distance of 5 cm from the UVC-LEDs (Corresponding to 0.245 mJ/cm^2^ and 0.065 mJ/cm^2^ per second, respectively; Fig. 2A). For both distances, the peak wavelength emission was measured at around 254 nm. For UVC-LED box 2, the emitted light intensity was measured with 117 µW/cm^2^ at 1 cm vertical distance from the LEDs, which corresponds to 0.117 mJ/cm^2^ per second (Fig. 2B). The peak wavelength emission was detected at 280 nm.

**Figure 1.**
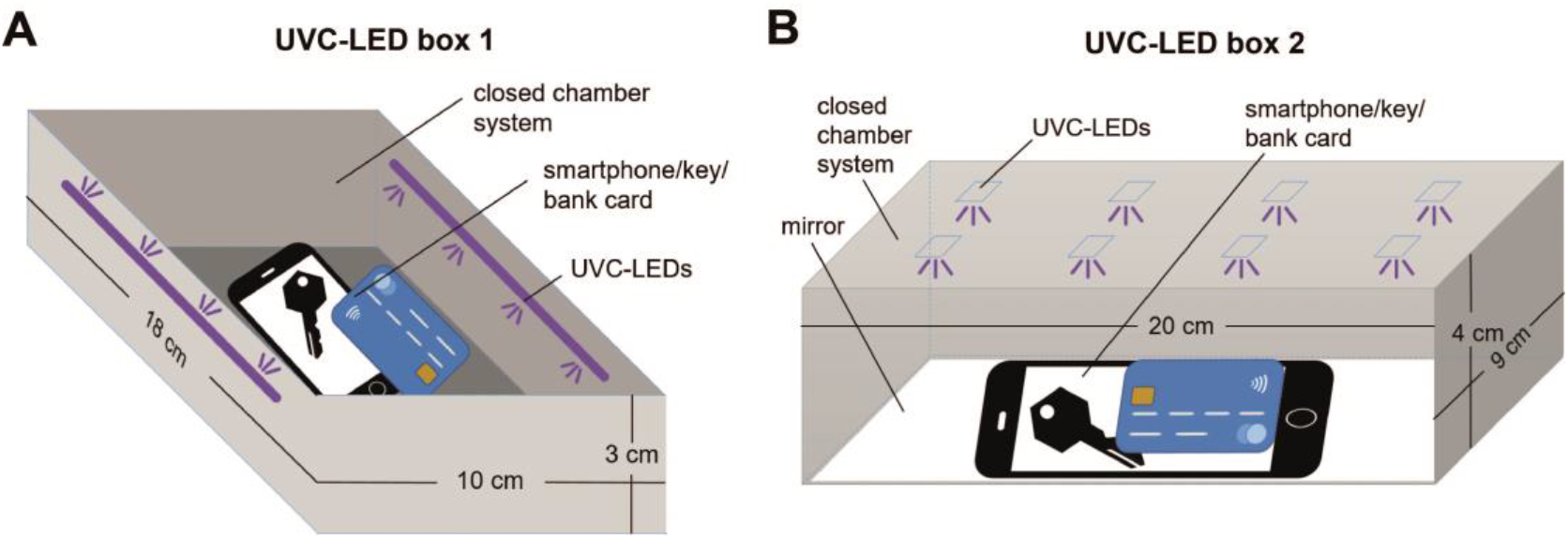
Schematic illustrations of the UVC-LED boxes used for UVC-disinfection. **A)** UVC-LED box 1 (Horcol) was equipped with lateral UVC-LEDs. **B)** UV-LED box 2 (expondo GmbH, Berlin, Germany) was equipped with UVC-LEDs incorporated in the lid and a mirror installed at the bottom of the chamber.

**Figure 2.**
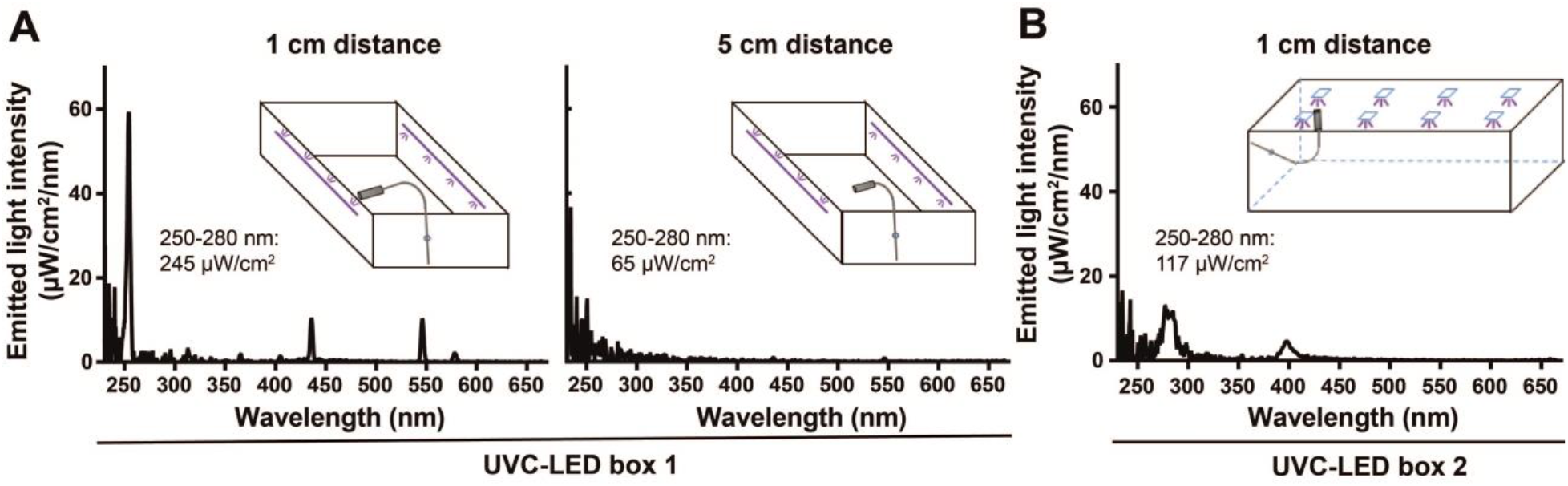
Spectrum of the emitted light intensity by LEDs inside of the UVC-LED boxes. The emitted light energy was measured by using a radiometrically calibrated spectrometer (STS-UV-L-50-400-SMA, Ocean Optics B.V., Ostfildern, Germany) with a sensitivity range between 190 and 650 nm (1.5 nm resolution) equipped with a CC-3-UV-S corrector (Ocean Optics B.V., Ostfildern, Germany). **A)** The spectrum of emitted light intensity of UVC-LED box 1 was measured at 1 and 5 cm horizontal distance from the lateral UVC-LEDs. **B)** The spectrum of emitted light intensity of UVC-LED box 2 was measured at 1 cm vertical distance from the UVC-LEDs. Emitted light intensity (µW/cm^2^) between 250 and 280 nm is displayed for each spectrum. Schematic illustrations indicate the borehole and the UVC-detector placed inside of the box for the respective measurement.

Next, metal, glass or plastic samples were overlaid with SARS-CoV-2 diluted in an organic matrix mimicking respiratory secretions. The final calculated virus concentration on the samples was 4.5 × 10^6^ TCID_50_/mL for UVC-LED box 1 and 1.8 × 10^6^ TCID_50_/mL for UVC-LED box 2, respectively. Compared to the viral load immediately after drying, there was no significant reduction of the viral loads after 10 minutes without irradiation (Fig. 3).

**Figure 3.**
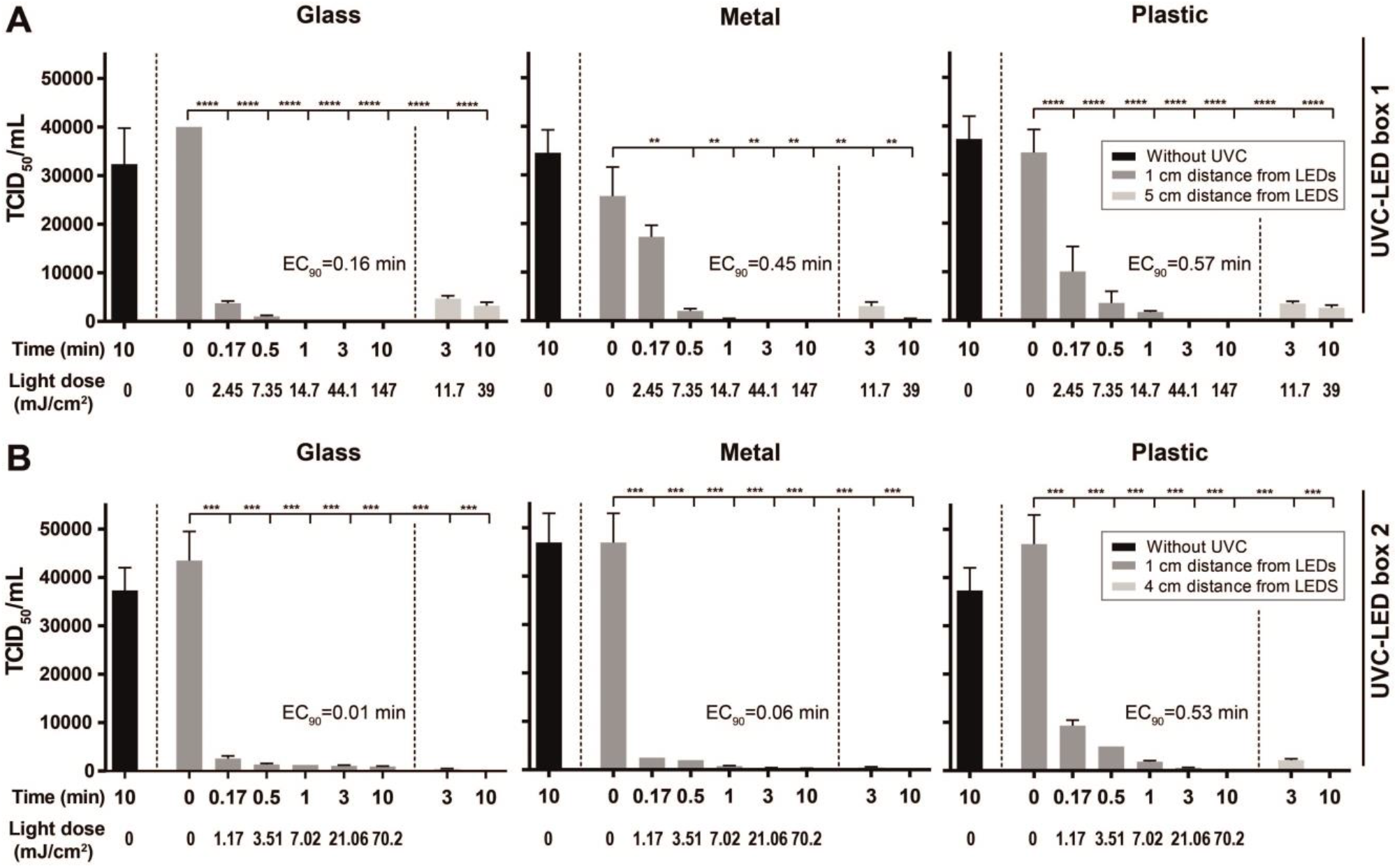
Disinfection of SARS-CoV-2 contaminated objects by two distinct UVC-LED boxes. Glass, metal and plastic samples were contaminated with SARS-CoV-2 at a final concentration of 4.5 × 10^6^ TCID_50_/mL for UVC-LED box 1 and 1.8 × 10^6^ TCID_50_/mL for UVC-LED box 2 in a cell culture medium mixed with defined organic matrix, thereby mimicking the viral contamination on the surfaces of personal belongings such as smartphones, keys, coins or credit cards. The samples were exposed to LED-UVC light for 0 s, 10 s, 30 s, 1 min, 3 min and 10 min at different distances from UVC-LEDs inside the UVC-LED boxes. Experiments were conducted in triplicates. A) In UVC-LED box 1, samples were irradiated at a distance of 1 cm and 5 cm from the UVC-LEDs. B) In UVC-LED box 2, samples were irradiated at 1 cm and 4 cm from the UVC-LEDs. Light doses at 4 cm distance cannot be shown, as an irradiance measurement of the LEDs at that distance would have required drilling a hole into the bottom of the box, which was not possible without damaging the box. Data are displayed as mean ± SD. *P < 0.05; **P < 0.01; and ***P < 0.001. TCID_50_=tissue culture infectious dose, 50%; EC_90_=90% effective concentration.

UVC-LED irradiation conducted with UVC-LED boxes proved to be an appropriate method for the disinfection of SARS-CoV-2 contaminated surfaces. SARS-CoV-2 contaminated glass, metal or plastic samples were effectively UVC-disinfected inside of both UVC-LED boxes (Fig. 3 and 4). A significant reduction of viral loads of SARS-CoV-2 on glass, metal and plastic was achieved even after 10 seconds of irradiation (UVC-LED box 1: 2.45 mJ/cm^2^; UVC-LED box 2: 1.17 mJ/cm^2^) at a distance of 1 cm from the LEDs in both UVC-LED boxes (Fig. 3). When using UVC-LED box 1, complete inactivation of SARS-CoV-2 could be achieved after 3 (glass and plastic) or 10 minutes (metal) of irradiation at a distance of 1 cm (Fig. 3A and 4A). At a distance of 5 cm from the LEDs (UVC-LED box 1), viral loads were strongly reduced after 3 and 10 minutes of irradiation (3 minutes: Glass: 88.27±1.49%, metal: 88.26±3.32%, plastic: 90±1.38%; 10 minutes: Glass: 91.92±1.85%, metal: 98.52±0.42%, plastic: 92.56±1.7%; Fig. 4A). When using UVC-LED box 2, SARS-CoV-2 was almost completely inactivated after 3 and 10 minutes of irradiation at a distance of 1 cm (3 minutes: Glass: 97.49±0.35%, metal: 99.21±0.1%, plastic: 98.74±0.16%; 10 minutes: Glass: 97.85±0.27%, metal: 99.42±0.08%, plastic: 99.46%; Fig. 4B). At a distance of 4 cm, SARS-CoV-2 was completely inactivated on glass and metal and almost completely on plastic (99.89±0.2%) after 10 minutes of irradiation (Fig. 3B and 4B).

**Figure 4.**
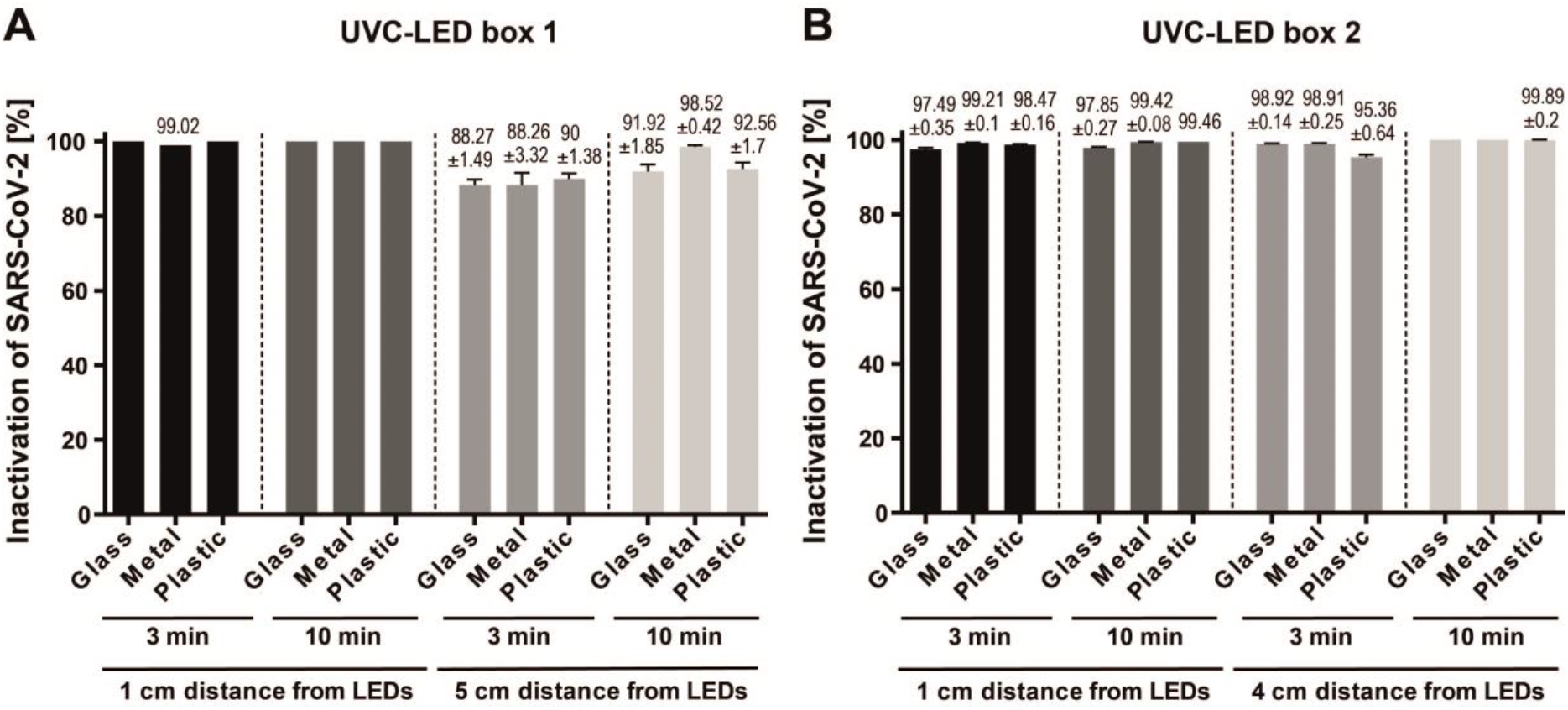
Percentage of inactivation of SARS-CoV-2 on surfaces after 3 or 10 minutes of UVC-LED irradiation. Glass, metal and plastic samples were contaminated with SARS-CoV-2 at a final concentration of 4.5 × 10^6^ TCID_50_/mL for UVC-LED box 1 and 1.8 × 10^6^ TCID_50_/mL for UVC-LED box 2 in cell culture medium mixed with defined organic matrix, thereby mimicking the viral contamination on the surfaces of personal belongings such as smartphones, keys, coins or credit cards. The samples were exposed to LED-UVC light for 0 s, 10 s, 30 s, 1 min, 3 min and 10 min at different distances from UVC-LEDs inside the UVC-LED boxes. Experiments were conducted in triplicates. A) In UVC-LED box 1, samples were irradiated at 1 and 5 cm from the UV-C LEDs. B) In UVC-LED box 2, samples were irradiated at 1 and 4 cm from the UV-C LEDs. Data are displayed as mean ± SD.

Taken together, our data demonstrate that UVC-LED sterilization boxes can effectively inactivate SARS-CoV-2 on surfaces such as glass, plastic or metal.

## Discussion

SARS-CoV-2 is transmitted through direct contact with infected individuals, by virus-containing aerosols and potentially via contact with virus-contaminated surfaces (*3*). In the present study, we investigated the performance of UVC-LED sterilization boxes on the inactivation of SARS-COV-2 on surfaces such as glass, metal and plastics that are typically found on personal items like smart phones, credit cards or keys. We demonstrated that UVC-LED boxes can effectively inactivate SARS-CoV-2 on glass, metal and plastic. Independent of the used UVC-LED box and the materials, SARS-CoV-2 could be almost completely inactivated after 3 minutes exposure (UVC-LED box 1: 1 cm: 44.1 mJ/cm^2^, 5 cm: 11.7 mJ/cm^2^; UVC-LED box 2: 1cm: 21.06 mJ/cm^2^), which represents the standard exposure time provided by the UVC-LED boxes. Even 10 seconds of exposure strongly reduced viral loads on the contaminated surfaces. The findings are in line with other studies that report susceptibility of coronaviruses, including SARS-CoV-1 and SARS-CoV-2 to UVC irradiation (*7, 10*). To almost completely inactivate high viral loads of SARS-CoV-1, a UVC-dose of 1446 mJ/cm^2^ was necessary (*10*). High viral loads of SARS-CoV-2 could be completely inactivated by a UVC dose of 1048 mJ/cm^2^ (*7*). The results of the present study show that the distance of the inoculated materials from the LEDs and thus the emitted light intensity is a decisive factor for achieving the complete inactivation of SARS-CoV-2. Accordingly, recently it has been reported that distance and angle of UVC light source in closed box systems are important factors for irradiance over respirator surfaces (*11*). UVC-LED box 1 and 2 showed a similar reduction of viral load for the materials glass and plastic. At a distance of 5 cm from the LED of UVC-LED box 1, although the viral load was reduced by 91.92±1.85% on glass, 98.52±0.42% on metal and 92.56±1.7% on plastic, the emitted light intensity (65 µW/cm^2^) was not sufficient to completely inactivate the virus after 10 minutes exposure. The mirror at the bottom of UVC-LED box 2 may have contributed to facilitate the inactivation of the virus. Van Doremalen et al. (*4*) indicated that a SARS-CoV-2 stock with a viral concentration of 10^5^ TCID_50_/mL corresponds to cycle-threshold values between 20 and 22, which is similar to the thresholds of samples from the upper and lower respiratory tracts of infected individuals. In this study, SARS-CoV-2 stocks with even higher viral concentrations of about 10^6^ TCID_50_/mL were used for the inoculation of the materials. Overall, both UVC-LED boxes were highly effective in inactivating the high titer viral stocks of SARS-CoV-2. These encouraging results make the UVC-LED boxes an affordable option for the public to disinfect a variety of items, including phones, watches, headphones, masks, makeup utensils, as long as the item size fits the device. As SARS-CoV-2 can also be detected on different surfaces in hospital environment (*12, 13*), UVC-LED boxes might also be an effective tool for environmental decontamination in hospitals.

## Acknowledgments

We thank Oliver Lischtschenko from Ocean Optics B.V. for providing us the spectrometer and his technical support.

This study was supported by the *Stiftung Universitätsmedizin Essen* (awarded to A. Krawczyk) and the Rudolf Ackermann Foundation (awarded to O. Witzke).

## Address for correspondence

Adalbert Krawczyk, Department of Infectious Diseases, West German Centre of Infectious Diseases, Universitätsmedizin Essen, University Duisburg-Essen, 45147 Essen, Germany; email

## Declaration of interest

The authors declare no conflict of interest.

## Notes

### Competing Interest Statement

The authors have declared no competing interest.

